# Allostery at a protein-protein interface harboring an intermolecular dynamic network

**DOI:** 10.1101/2024.01.07.574534

**Authors:** Sara Medina Gomez, Suresh Kumar Vasa, Rasmus Linser

## Abstract

Motional properties of individual amino acids in proteins are strongly modulated by their specific surrounding. The dynamics of tightly interacting residues can form *intra*molecular dynamic networks, which influence various features of protein function and serve as an access point for their modulation within signaling cascades. However, the possible formation of *inter*molecular networks shared between natural or constructed interaction partners has escaped thorough experimental assessment. Here, using fast-MAS solid-state NMR spectroscopy, we contrast the absence of a cross-talk between different residues in an apo protein with a recoupling of μs timescale dynamics effective via a mediating crystal-crystal contact. The data show that dynamic allostery is not necessarily restricted to motionally coupled elements within a single protein but can traverse molecular boundaries. Interrogation of intermolecular dynamic networks by the strategies proposed here may shed light on the mechanisms underlying allosteric modulation of protein function in biological, pharmacological, and biotechnological studies.

## Introduction

Adding to structure and site-specific chemical properties, motion constitutes a main pillar of protein functionality. In addition to translational motion and association/dissociation events, internal dynamics enable both, rather substantial changes between distinct conformations of vastly differential properties as well as more subtle local fluctuations that are equally important for the thermodynamic feasibility of such conformations, transitions, binding affinities, and chemical conversion.^1^ Such dynamics are inherently dependent on a complex array of internal and external parameters and hence allow the effective modulation of protein functionality as a function of upstream regulators, feedback loops, and other conditions.^2,3^

Owing to the physical connectivity between the individual moieties, creating a direct or indirect impact of one moiety on the motional properties of others, they form (*intra*molecular) dynamic networks with correlated motion, which govern the overall motional (and thermodynamic) personality of a protein.^4-6^ Orthosteric interaction partners, like small-molecule active-site ligands, can simply block interaction sites or access to functionalities, interfere with the chemical properties of active-site residues, or impact the effective shape of binding pockets locally. By contrast, dynamic networks, as complex arrangements of coupled elements, can also be detuned by *inter*molecular allosteric interactions.^1,7,8^ This impact can be derived from global shifts in conformational ensembles, e.g., stabilization of individual conformations out of a more heterogeneous energy landscape, upon inhibitor binding to sites spatially distant from the active site.^9-12^ Hence, binding-(in)competent or catalytically (un)productive conformations can be enriched.^13^ Whereas in some cases discrete conformations can be captured and the effect of a binder be rationalized, other cases are not associated with appreciable structural differences to the apo protein, and a more overarching impact onto the required protein dynamics has been assumed.^14^

NMR relaxation in solution and in solids has been used successfully to decipher motional timescales and the extent of dynamics for individual residues and collective motion in a large range of contexts.^15-19^ Nevertheless, it has remained hard to grasp how suitable dynamics can be unleashed by the association of binding partners. Also, pharmacological allosteric modulators mostly seem to hinge on tight interactions with their target.^12,20^ As such, it has remained unclear whether/how intermolecular dynamic networks at the interfaces between two or more *temporarily* interacting proteins can newly be formed. Our lack of understanding is due to the general difficulties for elucidation of the mechanical underpinnings of protein motion, but for this case in particular, it is further aggravated by the *transient* nature of binding for natural modulators, causing association/dissociation dynamics that technically interfere with the internal motional properties of interest.

Here we employ fast-magic-angle-spinning solid-state NMR to be able to tackle an intermolecular network of μs timescale dynamics at a protein-protein interaction site captured within a crystal, to circumvent any added association/dissociation dynamics. In particular, we use ^15^N relaxation dispersion of residues in the network to assess bond vector reorientation upon partial recoupling of anisotropic interactions close to the rotor-resonance condition in *R*_1π_ experiments. This framework, applied to residues at a crystal-crystal contact, serves to observe a motional coupling that is only induced upon intermolecular assembly of individual elements.

## Results

The RT loop in the monomeric SH3 domain of chicken α-spectrin in solution displays a rich spectrum of intrinsic flexibility.^21,22^ In previous work, we showed that in the context of a crystal lattice, some of its motional modes are slowed down to the μs regime, making it accessible to *R*_1π_ relaxation dispersion techniques in the framework of fast magic-angle spinning and proton detection.^21^ In contrast to ps-ns timescale motion in disordered loops (e.g., in the distal loop in SH3 domains, here residues 47, 48) and in most sidechains, comparison of experimental with simulated data for two-site exchange into different potential excited-state conformations suggested a backbone flip at the very tip of the RT loop as a statistically justified explanation.^21^ Fig. 1A shows the embedding of the central residue R21 in the crystal context, where the N- and C-terminal residues of a neighboring protein, which are only partly visible by diffraction methods, are revealed to be in close spatial proximity. In order to specifically probe the motional coupling of residue 21 with its intra- and intermolecular surroundings, we created a second subject of study by mutating residue R21 to alanine (See the SI for preparative details.) while leaving everything else identical. In the monomeric protein in solution, this mutation shows the expected behavior of a *local* perturbation and incurs major chemical-shift perturbations only at the very mutation site (Fig. 1B and C). The same is true for the replacement of R21 with aspartate, affording very similar CSPs. (See the SI, Fig. S1, for solution HSQCs and chemical-shift perturbations of the R21E mutant.) In the crystal, by much contrast, the R21A mutation entails CSPs in several areas of the protein primary sequence (Fig. 1D and E), with additional pronounced alterations at the N- and C-termini and around residues 31, 39, 49, and 57. (The R21E mutant did not crystallize in our hands.) Naively, these changes could simply be due to altered electrostatic effects sensed by multiple proximal residues at the monomer-monomer interface, even though they would be surprisingly widespread.

**Fig. 1:**
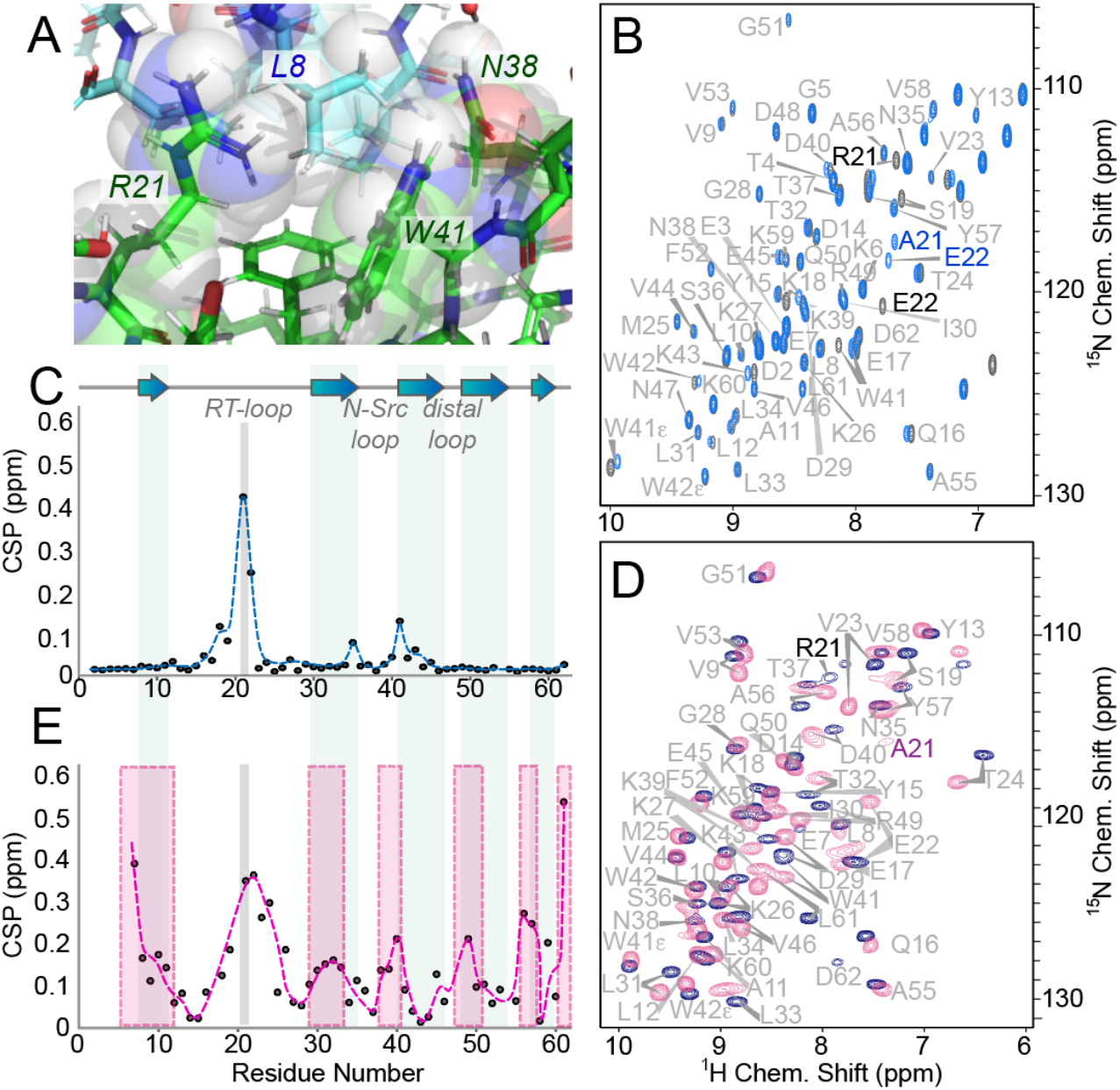
Steric context of the RT loop in a crystalline state and effects of its mutation. **A)** Zoom-in on the intermolecular interactions of the RT loop in SH3 crystals, using PDB structure 2NUZ. R21 and N38, as well as intermittent residues touching each other, are annotated. Green and blue structures denote different monomers. Van-der-Waals radii are depicted as semi-transparent. **B)** Overlay of HSQCs for wt (gray) and mutant (blue) SH3 in solution. **C)** Chemical-shift perturbations of the R21A mutation in solution. **D)** Overlay of solid-state NMR H/N correlations for wt (blue) and mutant (pink) SH3. **E)** Chemical-shift perturbations of the R21A mutation in the solid state. Regions affected on top of the local effect (gray line) are marked by pink boxes.

To interrogate the system in more detail, we looked closely at protein dynamics and their difference between the wt and mutant monomer. See Redfield relaxation data for ^15^N (*R*_1_, *R*_2_, heteronuclear steady-state NOE recorded at 298 K) as well as a reduced spectral-density mapping in Figs. S2 and 3. Fig. 2A depicts the *R*_2_ rates as a measure best representing contributions from both, fast and slow motions. In solution, we conclude that virtually no changes are introduced upon mutation, in particular not for the tip of the loop that carries the mutation. In the presence of the interactions within the crystal, on the other hand, motion at the mutation site is slightly increased. E. g., Fig. 2B shows that the *R*_1ρ_ rate at 20 kHz spin lock field strength rises from around 10.7 to 12.7 s^-1^. However, quite stunningly, dynamics here seem to be strongly affected also at the N- and C-terminus, N38, and the sidechain of W41 (Fig. 2B). In particular, *R*_1_ρ rates at 20 kHz spin lock field strength increase from 0.9 to 4.9, from 6.1 to 8.8, and from 5.6 to 12.0 s^-1^ for L61, W41:, and N38, respectively. N38 dynamics in the wt crystal have previously been observed to differ from the solution state.^23^ In addition, D62 and E7, with very high *R*_1_ρ rates already in the wt (7.7 and 16.3 s^-1^, respectively), are not visible/fittable at all anymore in the mutant crystal due to severely reduced signal-to-noise ratio. This signifies an even larger extent of motion in the mutant, now effectively evading dipolar magnetization transfers even more than for the wt. Altogether, a flexibilization seems to happen exactly for those residues participating in the network shown in Fig. 1A.

**Fig. 2:**
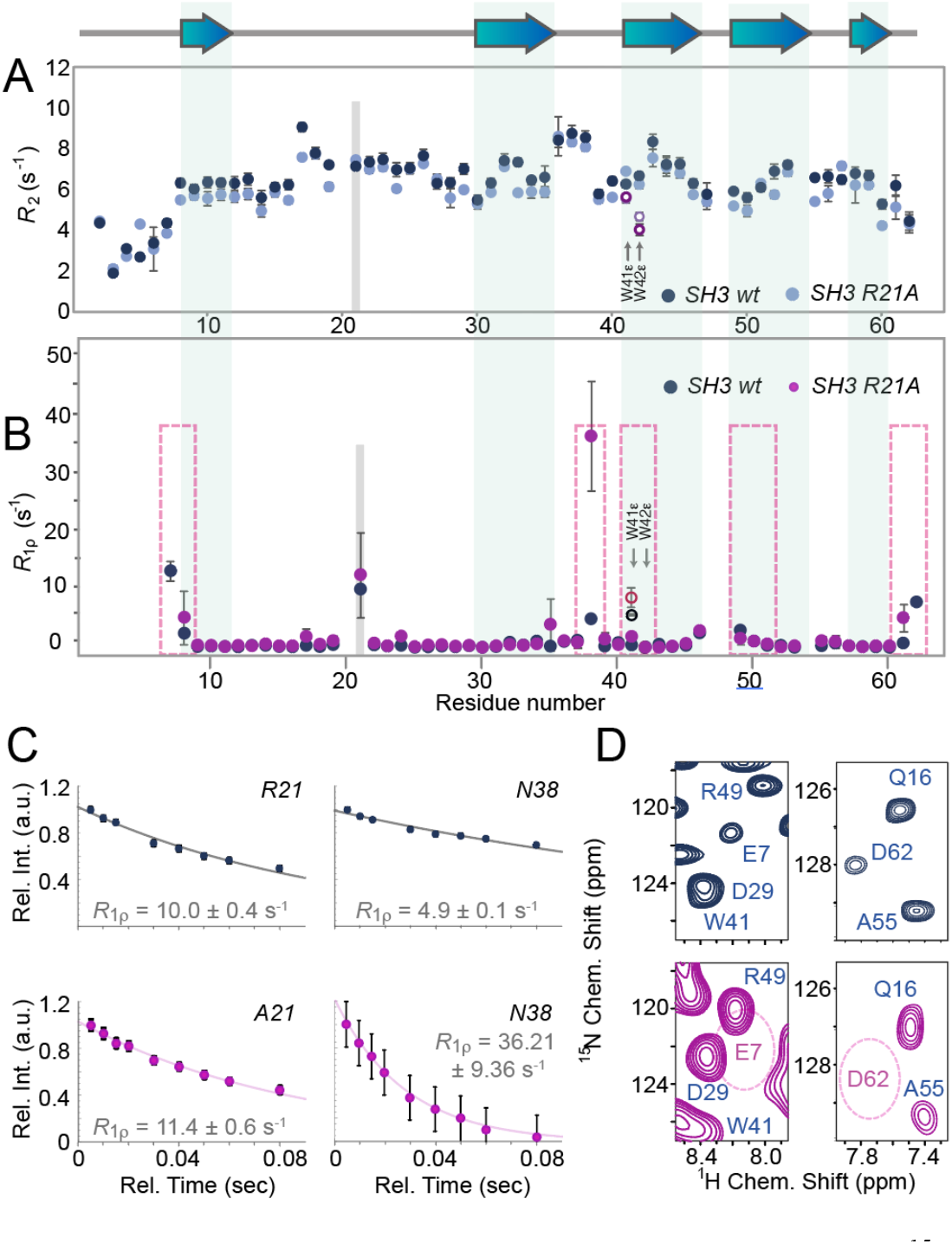
Local vs. global changes in the protein’s motional properties. **A)** Solution NMR ^15^N *R*_2_ rates as a function of sequence. **B)** Solid-state NMR ^15^N *R*_1ρ_ rates at 20 kHz spin lock field strength. **C)** ^15^N *R*_1_ρ decay profiles at 20 kHz for residues 21 (left) and 38 (right) in wt (top) and mutant (bottom). **D)** Weakening of E7 and D62 in wt and mutant protein.

In order to characterize the influence of the mutation onto the other spins in the neighborhood more closely, we conducted ^15^N near-rotary-resonance relaxation dispersion (NERRD) measurements for wt and R21A mutant in the solid state, for which a subset of data is shown in Fig. 3A. For rigid residues, the event of recoupling creates a sharp onset of strong relaxation at the rotor resonance condition. For residues moving on the μs timescale, however, the interference of internal motion and sample rotation creates a partial recoupling of anisotropic interactions even at spin lock field strengths significantly below and above the rotor-resonance condition.^24^ Position 21 shows the expected acceleration of its amide bond vector fluctuation induced by the shortening and “discharging” of the R21 sidechain. Now exceeding the typical timescale window of the NERRD measurements with an increasing broadening of the fast-motional behavior to a profile outside of the region of a standard fitting, the χ^2^-based significance criterion would suggest that no NERRD behavior can be reliably detected. Instead, a flat line, with a high y-axis offset (*R*_1ρ_^0^) is obtained as the best fit. Upon decreasing the temperature of the measurement (practically implemented by measuring in a smaller, 0.7 mm, rotor but maintaining the same spinning rate), the residue now completely escapes quantitative characterization, presumably due to a combination of the low CP transfer efficiency and exchange broadening (see Fig. S4). In any case, further increased dynamics for A21 compared to R21 in the wildtype can be deduced. More importantly, in agreement with the above, we see the same mobilization also for residues N38, L8, and L61 (the latter two representing the last residues of N- and C-terminus, respectively, that are still assessable in both constructs), as well as W41ε (compared in Fig. S5). (Fig. S6 shows *R*_1_ data, in which N38, L8, and W41ε are also increased, N38 changing by more than a factor of 2, *i. e*., from 0.7 to 1.8 s^-1^.) Fig. 3A also shows the NERRD profiles for N38 and L8. N38 again transitions from a nicely fittable profile into a similar situation as A21, and L8 and L61 are both “mobilized”, turning from profiles not far from the rigid-residue case to a prototypical NERRD behavior with pronounced μs timescale motion. (All profiles are shown in Figs. S8 and 9.) Fig. 3B displays the fitted order parameters of the individual residues as a function of sequence. In addition to the “starting values” for 1-S^2^ (for the wt protein before mutation), plotted on the X-ray structure in Fig. 3C, Fig. 3D shows the difference values between R21A and wt, displayed on the structure in the context of the interface.

**Fig. 3:**
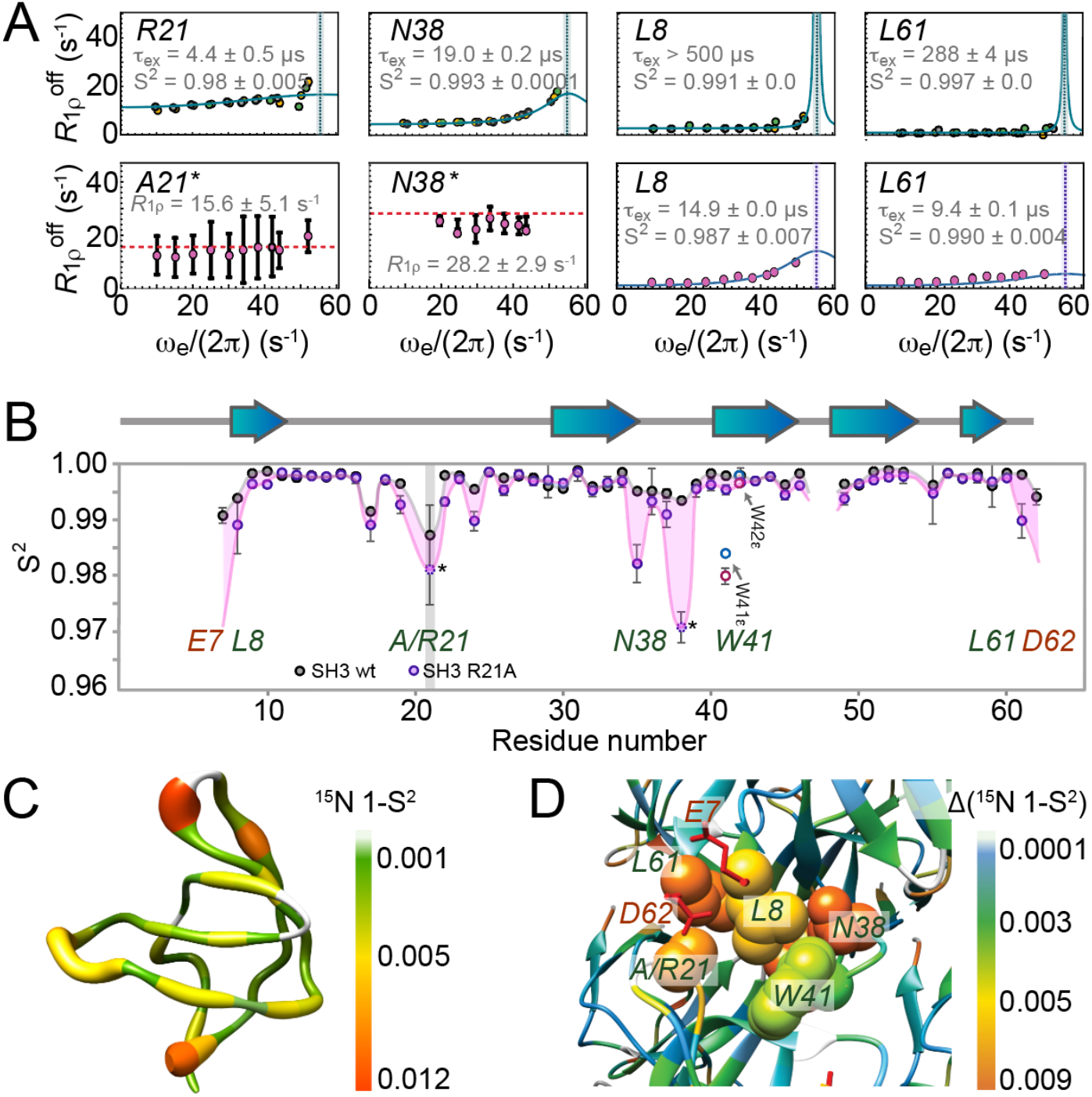
^15^N *R*_1_ρ NERRD data. **A)** NERRD profiles for residues 21, 38, 8, and 61 in the wt (top) and mutant protein (bottom). **B)** NERRD-derived order parameters as a function of sequence. For A21 and N38 in the case of the mutant (asterisks and dashed data points), NERRD fits were deliberately employed to extract μs timescale S^2^ even though their fast-motional behavior violates the chosen significance criterion compared to a flat line. The trendline for the termini of the mutant is extrapolated to reflect that reduced dipolar magnetization transfer efficiencies denote a further increase of motion. **C)** μs order parameters for SH3 wt depicted on the X-ray structure (2NUZ), representing NERRD-derived 1-S^2^ values both, as ribbon thickness and via the color code shown. **D)** Close-up of Δ(1-S^2^) between wt and mutant in the context of the crystal lattice. (X-Ray structure of the R21A mutant: PDB 2F2W; the W41 value accords to the N_ε_, which is more meaningful but almost identical to the backbone value, see C.)

The impression arises that the “flexibilization”, the increase of intrinsic dynamics, spans exactly those residues that are part of the network of physical contacts shown in Fig. 1A. In contrast to the solution case, their individual μs timescale motional properties seem to be coupled in the crystal. This also explains the strong chemical-shift perturbations in Fig. 1D/E, going beyond the local perturbations found for direct neighbors, even though the ground-state conformation (see an overlay of crystal structures in Fig. S7) seems unchanged upon mutation. Intriguingly, the motional dependency in focus seems to derive from intermolecular interactions brought upon by the crystal-crystal contact, which brings the N- and C-terminal residues in to “fill up” a vacancy of direct interactions between positions 21 and 38 otherwise. We suggest the following mechanistic explanation: In the event of μs timescale motion at the tip of the RT loop, assuming, e. g., the previously suggested peptide flip^21^, a conformational exchange that is also seen as a mix of crystallographic ground state structures upon R21G mutation,^22^ or simply a slow sidechain reordering of R21, a concomitant reordering of the vicinal sidechains would occur. For the shorter residue 21 sidechain in the mutant, this dynamics is associated with lower steric restraints, which makes it faster. At the same time, the steric constraints for vicinal sidechains are relaxed, and their possibilities to find the respective lowest-energy conformation are improved, allowing larger-amplitude changes within the network of sterically coupled residues and hence faster and larger-amplitude motion up to residue N38. Strictly speaking, the details of the mutual interactions could be slightly different, e. g., additionally including other backbone and sidechain atoms. Nevertheless, the data strongly suggest that the structures at this protein interface establish an intermolecular dynamic network derived from loose, steric interactions that is non-existing in the monomeric protein, where the chain of sequential contacts between positions 21 and 38 is not applicable.

The experimental demonstration of intermolecular protein-protein interactions that *induce* coupling of μs timescale motion between residues of the same protein sheds light on how biological interaction partners can allosterically influence protein functionality without firm, visible changes in ground-state structures. In particular, the experiment demonstrates why allosteric modulation cannot only modulate target function destructively due to disruption of local dynamics or of dynamic networks, e. g., through steric blockage or rigidification of protein segments, as associated with some orthosteric inhibitors^25,26^. Instead, allosteric modulation seems possible in a constructive way, by adding missing links between individual residues or into partially preformed networks. The data show that interaction partners can fill in otherwise missing nodes to generate dynamic coupling, thereby significantly altering the landscape of conformational entropy. Such changes will hence affect binding/selectivity properties or enzymatic activity of the protein. Whereas such constructive mechanisms are less likely for small-molecule allosteric modulators, the picture obtained here may be representative for manifold regulatory protein-protein interactions, in which the intermolecular alteration of dynamics can tailor context-specific protein functionality.

Gaining experimental access to long-range motional coupling induced via protein-protein interactions by solid-state NMR spectroscopy generally represents itself as an attractive approach to study biological conglomerates that are too short-lived for assessment of relaxation (dispersion) studies in solution. The existence of additional crystal-crystal contacts other than the interaction surfaces in focus compared to a pure heterodimer in solution can potentially distort the picture. Nevertheless, we think that qualitative/mechanistic information on the interplay at the surface will provide important insights and possibly inform on further experimental routes for an improved understanding of the research target. Beyond the described protein-protein interactions, DNA, RNA, or even non-biological surfaces, which can equally trigger protein functionality through dynamic allostery, are possible extensions to the suggested methodology.

In conclusion, using ^15^N relaxation and relaxation dispersion behavior in solution compared to solid-state NMR of protein crystal-crystal contact, we demonstrated the interplay of amino acids from different monomers to constitute a transductive element generating an intermolecular dynamic network. Contrasting the absence of motional changes upon mutation of a loop residue in the monomeric protein, the mutation entails stark changes within a group of residues from two different monomers when these are brought into close proximity in the solid state. The experiments elucidate that motional coupling can be switched on via protein-protein association, demonstrating a possible mechanism of allosteric regulation via newly created intermolecular networks.

## Acknowledgments

Funded by the Deutsche Forschungsgemeinschaft (DFG, German Research Foundation) under Germany’s Excellence Strategy -EXC 2033 – 390677874 – RESOLV, and EXC-114 – 24286268 – CiPS-M. Funded by the Deutsche Forschungsgemeinschaft (DFG, German Research Foundation) – 27112786, 325871075 and the Emmy Noether program.

